# Granulocyte Colony Stimulating Factor causes cerebellar deficits and anxiety in a mouse model of CSF-1 receptor-related leukoencephalopathy

**DOI:** 10.1101/2022.02.21.481325

**Authors:** Fabrizio Biundo, Violeta Chitu, Jaafar Tindi, Nesha S. Burghardt, Gabriel G. L. Shlager, Harmony C. Ketchum, Michael A. DeTure, Dennis W. Dickson, Zbignew K. Wszolek, Kamran Khodakhah, E. Richard Stanley

## Abstract

Colony stimulating factor (CSF) receptor-1 (CSF-1R)-related leukoencephalopathy (CRL) is an adult-onset, demyelinating neurodegenerative disease caused by autosomal dominant mutations in *CSF1R,* modeled by the *Csf1r^+/-^* mouse. The expression of *Csf2,* encoding granulocyte-macrophage CSF (GM-CSF) and of *Csf3,* encoding granulocyte CSF (G-CSF), are elevated in both mouse and human CRL brains. While monoallelic targeting of *Csf2* has been shown to attenuate many behavioral and histological deficits of mouse CRL, including cognitive dysfunction and demyelination, the contribution of *Csf3* has not been explored. In this manuscript, we investigate the behavioral, electrophysiological and histopathological phenotypes of CRL mice following monoallelic targeting of *Csf3.* We show that *Csf3* heterozygosity normalized the *Csf3* levels in *Csf1r^+/-^* mouse brains and ameliorated anxiety-like behavior, motor coordination and social interaction deficits, but not their cognitive impairment. Consistent with this, *Csf3* heterozygosity attenuated microglial activation in the cerebellum and in the ventral but not in the dorsal hippocampus. *Csf3* heterozygosity also failed to prevent demyelination. *Csf1r^+/-^* mice exhibited altered synaptic activity in the deep cerebellar nuclei (DCN) associated with increased deposition of the complement factor C1q on glutamatergic synapses and with increased engulfment of glutamatergic synapses by DCN microglia. These phenotypes were significantly ameliorated by monoallelic deletion of *Csf3*. Our findings indicate that G-CSF and GM-CSF play non-overlapping roles in mouse CRL development and suggest that G-CSF could be an additional therapeutic target in CRL.

## Introduction

*CSF1R*-related leukoencephalopathy (CRL), previously named adult-onset leukoencephalopathy with axonal spheroids and pigmented glia (ALSP), pigmentary orthochromatic leukodystrophy (POLD) and hereditary diffuse leukoencephalopathy with spheroids (HDLS), is a neurodegenerative disease characterized by progressive cognitive impairment, motor coordination deficits, and psychiatric symptoms (1, 2). CRL is caused by autosomal dominant mutations in the colony stimulating factor-1 receptor gene *(CSF1R)* that inhibit the kinase activity or abolish the expression of the mutant chain (3). Based on the finding that haploinsufficiency is sufficient for the development of CRL in humans (4), we have validated the *Csf1r^+/-^* mouse as a model of CRL that reproduces the neurocognitive deficits and histopathological features of the human disease (reviewed in (5)). Quantitative transcriptomic profiling of autopsied brain samples from patients with CRL revealed the loss of homeostatic microglia suggesting that CRL might be a primary microgliopathy (6, 7). This concept has been reinforced by studies in the mouse CRL model showing that *Csf1r* heterozygosity in microglia was sufficient to reproduce all aspects of disease (8). In a screen for inflammatory cytokines, chemokines and receptors that could contribute to disease we found that the mRNAs for *Csf2,* encoding granulocyte-macrophage CSF (GM-CSF) and for *Csf3,* encoding granulocyte CSF (G-CSF), were uniquely elevated in the brains of *Csf1r^+/-^* mice (9). Elevation of *CSF2* expression in CRL patient brains (10) and the identification of gene expression changes consistent with altered G-CSF signaling (6, 7) suggested that both GM-CSF and G-CSF may also contribute to the development of this disease. Notably, while the expression of transcripts for both growth factors is barely detectable in normal brains, they can be rapidly induced by a variety of inflammatory stimuli, tissue injury and neurotoxins and signal in microglia to promote functional changes (reviewed in (11)). GM-CSF is a microglial mitogen (12, 13) that promotes a demyelinating phenotype in microglia (14), while G-CSF induces a pro-oxidant phenotype (15).

Genetic targeting in the mouse CRL model revealed that *Csf2* was responsible for the cognitive and olfactory deficits of *Csf1r^+/-^* mice and for callosal demyelination and atrophy (10). Furthermore, gene expression profiling of isolated forebrain microglia revealed maladaptive functions of *Csf1r^+/-^* microglia and activation of pathways triggering oxidative stress that were relieved by monoallelic *Csf2* deletion (10). However, although targeting *Csf2* improved coordination on the balance beam, it did not resolve the ataxic behavior in female mice or cerebellar microgliosis (10).

In the present study, we have explored the role of G-CSF in CRL pathology. We show that *CSF3* mRNA is also elevated in CRL patient brains and that the elevation of *Csf3* mRNA in the *Csf1r^+/-^* CRL mouse can be normalized by monoallelic *Csf3* deletion. In contrast to *Csf2* heterozygosity, monoallelic targeting of *Csf3* failed to prevent the cognitive deficits, callosal microgliosis and demyelination in the brains of CRL mice. However, *Csf3* heterozygosity attenuated the anxiety-like behavior, motor coordination and social novelty deficits of CRL mice. Consistent with these effects, monoallelic targeting of *Csf3* reduced microglial activation in cerebellum and ventral hippocampus, two brain regions involved in motor coordination and anxiety, respectively (16, 17). *Csf1r^+/-^* mice exhibited altered electrophysiological responses in the deep cerebellar nuclei (DCN) that were associated with increased expression and deposition of the C1q factor of the complement cascade on glutamatergic synapses and their increased engulfment by DCN microglia. All these phenotypes were attenuated in *Csf1r^+/-^; Csf3^+/^*^-^ mice. Together, our data suggest that in CRL, increased G-CSF promotes anxiety and cerebellar dysfunction by activating discrete populations of microglia and acts in a non-overlapping manner with GM-CSF to promulgate the disease.

## Results

### *Csf3* expression is elevated in mouse and human CRL brains and normalized in mice by *Csf3* heterozygosity

In a previous study, we observed an increased expression of *Csf3* in brains of pre-symptomatic *Csf1r^+/-^* CRL mice. The elevation of *Csf3* became more pronounced in aged mice exhibiting behavioral deficits suggesting a role for G-CSF in the CRL phenotype (9). To investigate whether *CSF3* expression was increased in CRL patients, we quantified the levels of *CSF3* mRNA in brains of CRL patients by qPCR. Consistent with the results detected in the mouse model, the levels of *CSF3* transcripts were significantly higher in CRL than in control brains (Fig. 1A). These results prompted us to generate and characterize an experimental cohort of mice including mice in which either one allele of *Csf1r* or of *Csf3* was deleted (*Csf1r^+/-^*, *Csf3^+/-^*), double mutants (*Csf1r^+/-^* ;*Csf3^+/-^*, referred to as *Dhet*) and wild type (*wt*) controls. Measurement of *Csf3* expression showed that the elevated *Csf3* mRNA levels observed in *Csf1r^+/-^* mice were normalized by *Csf3* monoallelic deletion (Fig. 1B).

**Figure 1.**
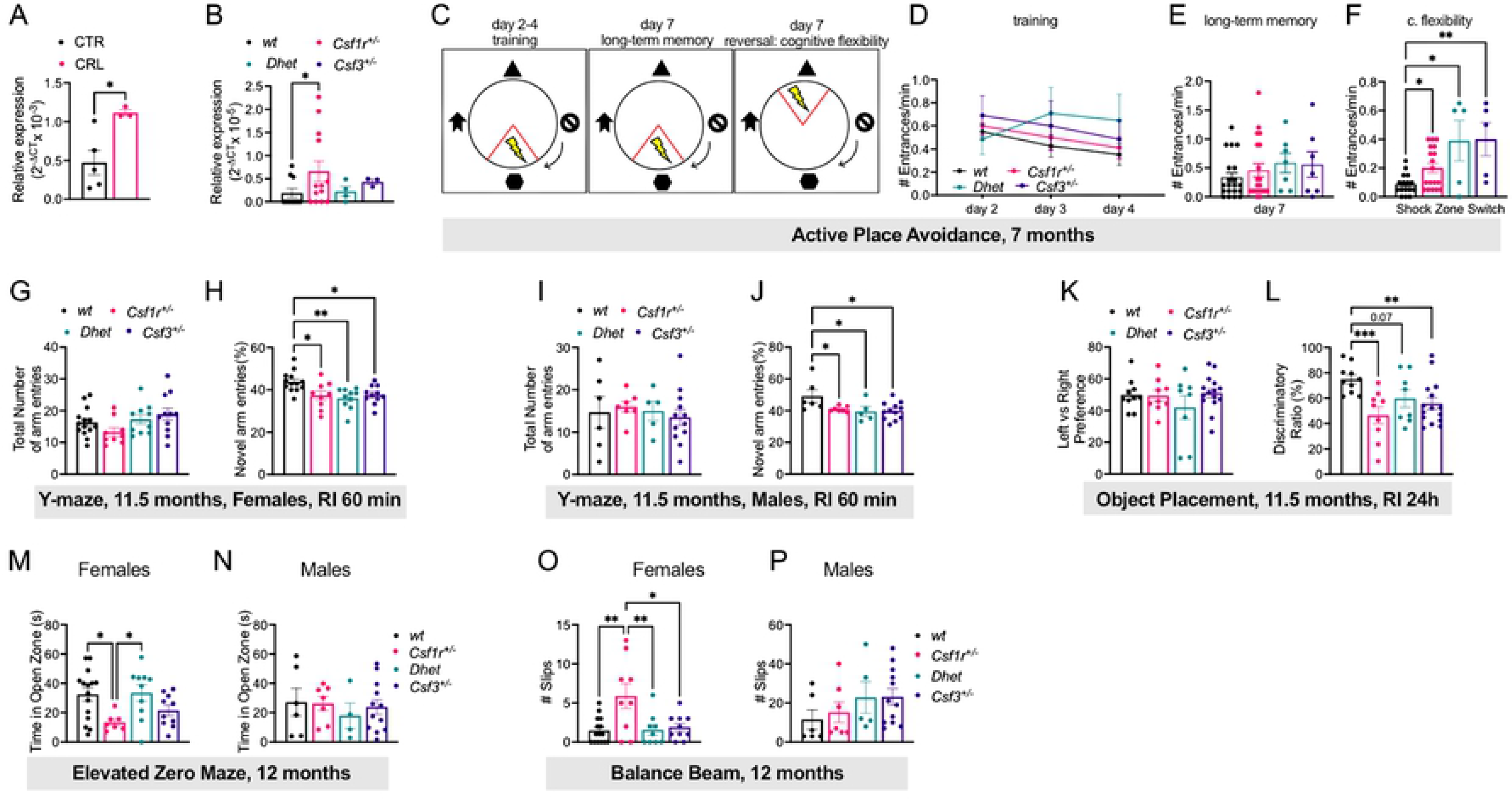
*Csf3* heterozygosity attenuates anxiety and motor coordination deficits in CRL mice but fails to improve cognition. **(A)** Elevated expression of *CSF3* in CRL patients versus healthy controls (unpaired t test). **(B)** Expression of *Csf3* in brains of *wt* and mutant mice (unpaired t test). **(C-F)** Evaluation of cognitive flexibility in 7- month-old mice (females plus males). **(C)** Schematic of the protocol used for active place avoidance testing. Day 1 (habituation) is not shown. **(D)** Days 2-4: Training to avoid the initial shock zone location. **(E, F)** Evaluation of long-term memory three days after the last training trial **(E).** Evaluation of cognitive flexibility after the location of the shock zone was switched **(F)** at day 7 (uncorrected Dunn’s test). **(G-J)** Assessment of short-term memory at 11.5 months of age in the Y-maze. **(G, H),** females; **(I, J)**, males. **(G, I)** Comparable total exploratory activity among genotypes. **(H, J)** Exploratory preference for the novel arm (Tukey’s). **(K, L)** Assessment of long-term memory in the object placement test (females plus males). **(K)** Ratio of the time exploring the left vs right position of the objects during training. **(L)** Discriminatory ratio of the time exploring the displaced vs the non-displaced object during testing (Fisher’s). **(M, N)** Assessment of anxiety-like behavior for females **(M)** and males **(N)** in the elevated zero maze (Bonferroni’s). **(O, P)** Measurement of motor coordination on the balance beam (Fisher’s). Means ± SEM, significantly different changes are marked by asterisks. *, p < 0.05; **, p < 0.01; ***, p < 0.001, ****, p < 0.0001. The statistical test used in each panel is indicated in parenthesis in the corresponding description. RI, retention interval. Data underlying this figure can be found in Table S1.

**Table 1:**
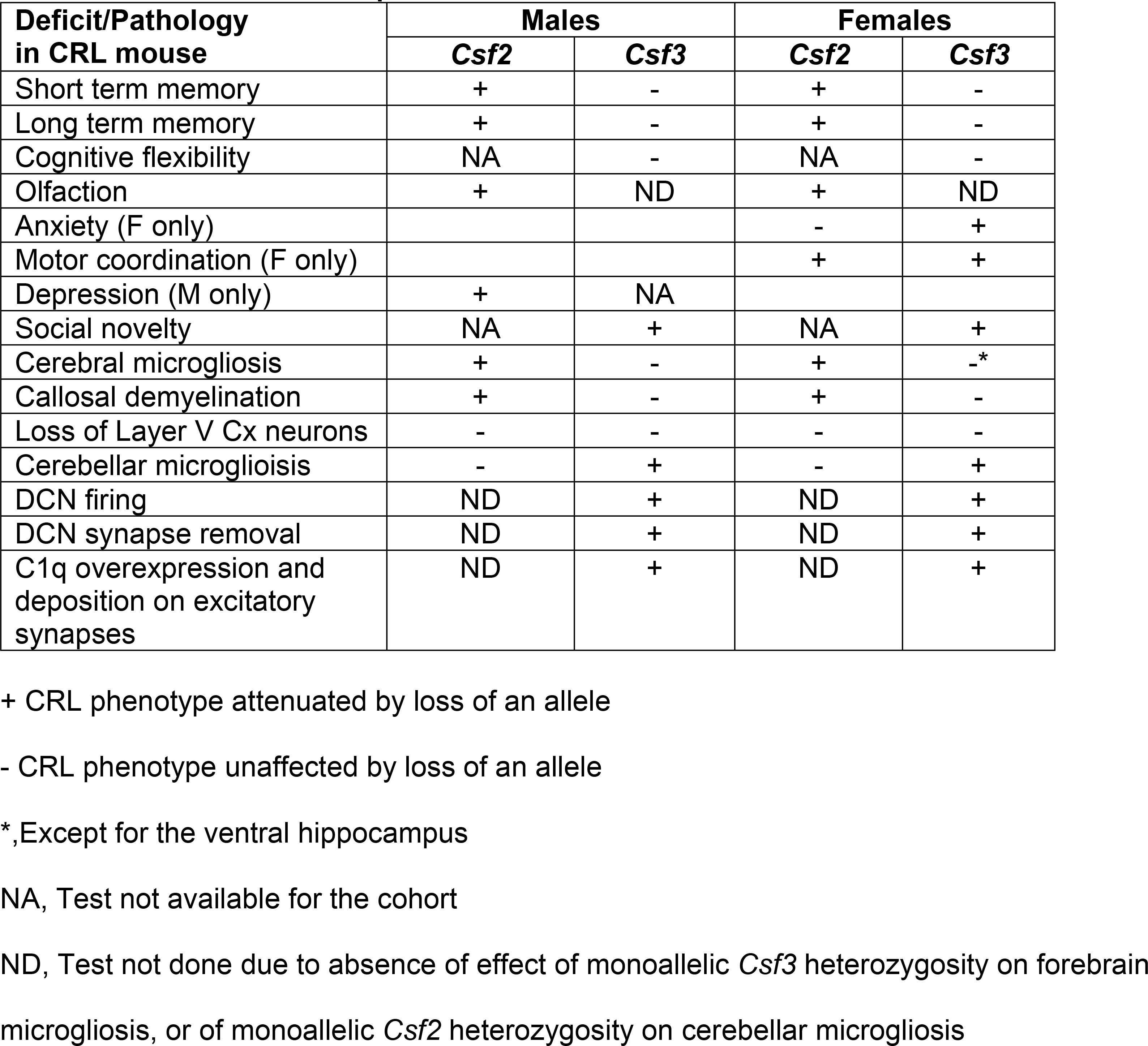
Effects of elevated Csf3 and Csf2 in mouse CRL illustrating their complementary contribution to disease development

### *Csf3* heterozygosity fails to prevent the cognitive deficits of CRL mice but attenuates anxiety-like behavior and the motor coordination deficits

To assess the contribution of the increased *Csf3* expression to the behavioral deficits of *Csf1r^+/-^* mice, the experimental cohort was evaluated for cognitive flexibility, spatial memory, anxiety, and motor coordination.

Cognitive flexibility, defined as the ability to change and adapt behavior in response to new environmental stimuli (18), is one of the executive functions affected in early stages of Alzheimer’s disease (19) and deficits were also recently reported in a case of CRL (20). Cognitive flexibility was evaluated in 7-month-old mice using the active place avoidance test (Fig. 1C) (21). Regardless of genotype, all mice had the same propensity to avoid the shock zone three days after the last training trial (Fig. 1D, E). These data indicate that at this young age, there are no significant long-term memory deficits associated with *Csf1r* and/or *Csf3* heterozygosity. However, when the location of the shock zone was switched, all mice carrying mutations entered the new shock zone significantly more than wt mice, demonstrating that either single or combined *Csf1r* and *Csf3* deficiencies impair cognitive flexibly (Fig. 1F).

The effects of *Csf3* heterozygosity on short- and long-term spatial memory were tested in aged (11.5 month-old) mice, in the Y-maze and object placement tasks, respectively (Fig. 1G-M).

All mutant mice (*Csf1r^+/-^*, *Csf3^+/-^* and *Dhet)* exhibited a short-term memory deficit as shown by their loss of preference for the novel arm of the Y-maze (Fig. 1H, J). The absence of differences in total number of arm entries indicated that the differences in cognitive performance did not result from different propensities among groups to explore of the apparatus (Fig. 1G, I).

Similarly, in the object placement test, all mutant mice showed no preference towards exploring the displaced object versus the non-displaced object, indicative of a long-term memory deficit (Fig. 1L). The absence of differences in time exploring the two initial positions of the objects during training indicated that the cognitive deficits detected during testing were not due to a preferential exploration of one of the sides of the chamber (Fig. 1K).

Mice were tested for anxiety-like behavior in the elevated zero maze at the age of 12 months (Fig. 1M, N). The time spent in the open zone of the circular apparatus was used as an index inversely related to anxiety-like behavior. Deficits observed in female *Csf1r^+/-^* mice (Fig. 1M) were prevented by *Csf3* heterozygosity, while no significant differences among the genotypes were detected in males ((Fig. 1N).

Motor coordination was analyzed at the age of 12 months using the balance beam test (Fig. 1O, P). The deficits observed in female *Csf1r^+/-^* mice were rescued by single-allele deletion of *Csf3* (Fig. 1O), while no significant differences were observed in males (Fig. 1P). Thus, *Csf3* heterozygosity fails to prevent the cognitive deficits of *Csf1r^+/-^* mice, but significantly attenuates the anxiety and loss of motor coordination observed in females.

### *Csf3* heterozygosity reduces microglial activation in the ventral but not dorsal hippocampus of CRL mice

The dorsal hippocampus plays a critical role in cognition while the ventral hippocampus relates to emotions and stress (22). Previous studies have shown that dysregulation of *Csf1r* signaling in microglia of CRL mice leads to low grade microgliosis in multiple regions of the brain, including the dorsal hippocampus (8, 10) and administration of recombinant G-CSF was reported to increase the number of microglia and their activation *in vivo* (15, 23). To test whether G-CSF regulates microglial activation in the hippocampus, we analyzed microglia densities and morphology in 16-month-old mice (Fig. 2). Iba1 staining revealed that microglial densities were significantly increased in the dorsal hippocampi of *Csf1r^+/-^* mice, and that *Csf3* heterozygosity failed to attenuate this increase (Fig. 2A - upper panels and Fig. 2B), yet there were no significant differences in microglial densities in the ventral hippocampus (Fig. 2A - lower panels, Fig. 2C). Interestingly, while the branching and length of microglial processes did not vary with genotype in the dorsal hippocampus (Fig. 2D – right panels, Fig. 2E), both the branching and length of microglial processes were decreased in the ventral hippocampus of *Csf1r^+/-^* mice (Fig. 2D – left panels, Fig. 2F), indicative of an activated state. Monoallelic deletion of *Csf3* restored process branching, albeit it had no significant effect on process length (Fig. 2F). These data are consistent with a contribution of G-CSF-induced microglial activation to the development of anxiety-like behavior, but not to the cognitive deficits in *Csf1r^+/-^* mice (Fig. 1C-M).

**Figure 2.**
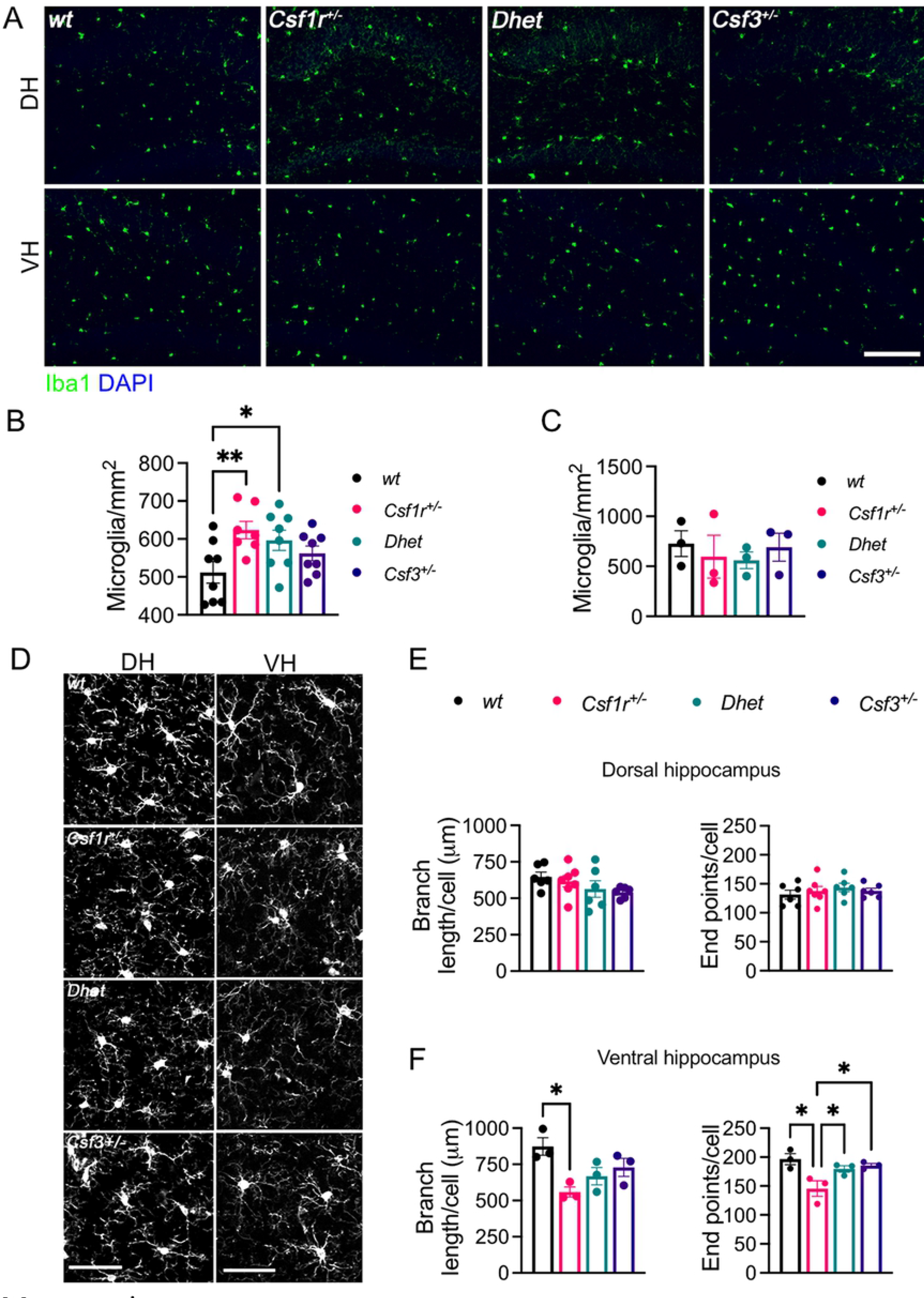
*Csf3* heterozygosity reduces microglial activation in the ventral hippocampus of CRL mice. **(A)** Iba1^+^ cell densities (green) in dorsal hippocampus (DH) and ventral hippocampus (VH) of 16- month-old female mice (scale bar, 100 μm). **(B, C)** Quantification of microglial densities in the DH (B) and VH (C), 3-8 mice per genotype) (Fisher’s). **(D)** Morphology of the microglial ramifications in the dorsal and ventral hippocampus (scale bar, 50 μm). **(E, F)** Quantitation of the ramifications in DH **(E)** and VH **(F)**, 3-6 mice per genotype. (Two-stage linear step-up procedure of Benjamini, Krieger and Yekutieli). Means ± SEM, significantly different changes are marked by asterisks. *, p < 0.05; **, p < 0.01; ****, p < 0.0001. Data underlying this figure can be found in Table S1.

### *Csf3* heterozygosity fails to prevent microglial activation, callosal demyelination and neurodegeneration in the motor cortex of CRL mice

Previous studies have shown that dysregulation of CSF-1R signaling in microglia of CRL mice promotes callosal demyelination and the loss of layer V neurons in the motor cortex (10). To test whether G-CSF plays a role in microglial activation in the corpus callosum, we analyzed microglia density and morphology in 16-month-old CRL and *wt* control mice. Consistent with the previously published data (10), analysis of multiple sagittal brain sections detected a significant elevation in the total number of microglia and in the number of sections with supraventricular microglial patches in the corpus callosum of *Csf1r^+/-^* compared to *wt* mice (Fig. 3A, B). Although *Csf3* heterozygosity reduced the total number of microglia, it did not significantly reduce their propensity to cluster (Fig. 3B, right panel) and failed to prevent the shortening of their processes and the loss of process branching (Fig. 3C, D). The presence of clusters of activated microglia has recently been correlated with the active clearing of degenerated myelin (24). Consonant with its inability to reduce microglial clustering and activation, *Csf3* heterozygosity also failed to attenuate callosal demyelination (Fig. 3E, F). Furthermore, histological evaluation of the motor cortex revealed that although *Csf3* heterozygosity attenuated the shortening of cortical microglia processes, it failed to attenuate their expansion in the motor cortex (Fig. 3 G-J). Consistent with this, *Csf3* heterozygosity also failed to prevent the loss of Layer V neurons (Fig. 3K, L). These data indicate that, although G-CSF may regulate some aspects of callosal and cortical microglia activation, its actions do not contribute significantly to callosal demyelination or to neurodegeneration in the motor cortex.

**Figure 3.**
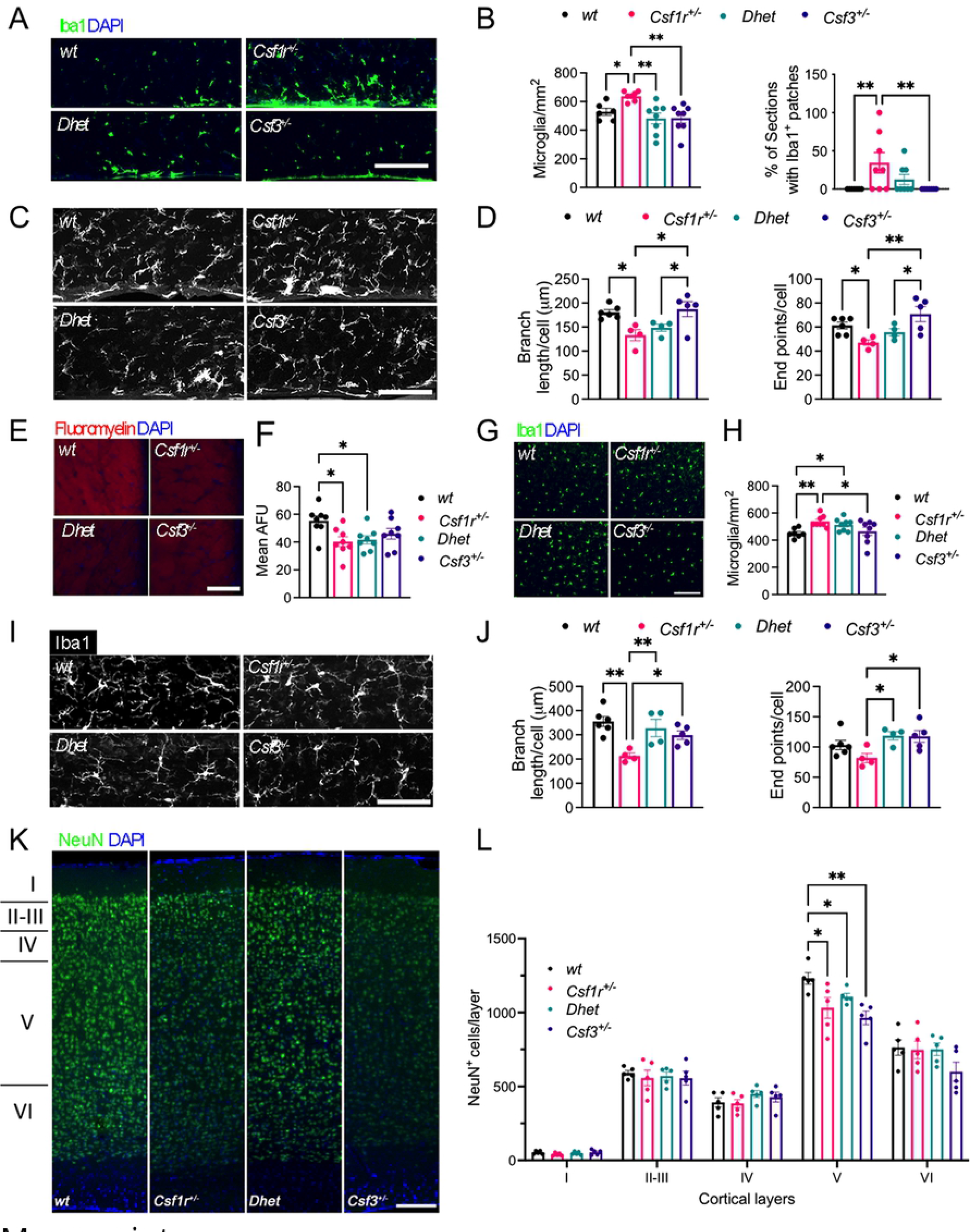
*Csf3* heterozygosity does not prevent microglial activation in the corpus callosum or motor cortex, nor callosal demyelination or cortical neuronal loss, in CRL mice. **(A)** Iba1^+^ cell densities (green) in the supraventricular area of corpus callosum (scale bar, 100 μm). **(B)** Quantification of microglial densities (left panel) and percentage of sections with microglial patches (right panel) (6-8 mice per genotype) (Fisher’s). Morphology **(C)** and morphometric analysis **(D)** of microglia in corpus callosum (scale bar, 50 μm) (4-6 mice per genotype) (Two-stage linear step-up procedure of Benjamini, Krieger and Yekutieli). Fluoromyelin staining **(E)** and fluorescence quantification **(F)** in corpus callosum (scale bar, 100 μm) (6-8 mice per genotype) (Tukey’s). Iba1^+^ cell densities (green) **(G)** and quantification **(H)** in the motor cortex (scale bar, 100 μm) (7-8 mice per genotype) (Fisher’s). Morphology **(I)** and morphometry **(J)** of microglia in the motor cortex (scale bar, 50 μm) (4-6 mice per genotype) (Two-stage linear step-up procedure of Benjamini, Krieger and Yekutieli). NeuN^+^ neurons **(K)** and quantification **(L)** of their distribution in the cortical layers of the motor cortex (scale bar, 100 μm) (5 mice per genotype) (Fisher’s). All experiments were performed in 16-month-old female mice. Means ± SEM, significantly different changes are marked by asterisks. *, p < 0.05; **, p < 0.01. Data underlying this figure can be found in Table S1.

### *Csf3* heterozygosity prevents microglial activation in the cerebellum

In addition to the corpus callosum and the motor cortex, the cerebellum plays an important role in motor coordination. This prompted us to analyze the impact of *Csf3* heterozygosity on microglia density and activation in the cerebellum. Analysis of the cerebellar cortex and deep cerebellar nuclei revealed an increase in the number of Iba1^+^ microglial cells detected in the cerebellar cortex of *Csf1r^+/-^* mice compared to *wt* mice. This was attenuated by monoallelic deletion of *Csf3* (Fig. 4A, B). In contrast, no differences in microglia densities were detected in the dorsal protuberance of the medial cerebellar nucleus (MedDL), and the interposed cerebellar nuclei (Int) (Fig. 4A, B). Morphometric analysis revealed an increase in microglia activation in *Csf1r^+/-^* mice in all the areas of the cerebellum that was prevented by *Csf3* monoallelic deletion (Fig. 4C-right panels and Fig. 4D, F-H). Furthermore, *Csf3* heterozygosity also reduced the extent of microglia contacts with the Purkinje cell somas (Fig. 4C- left panels and Fig. 4E). These data indicate that G-CSF mediates the activation of cerebellar microglia in *Csf1r^+/-^* mice.

**Figure 4.**
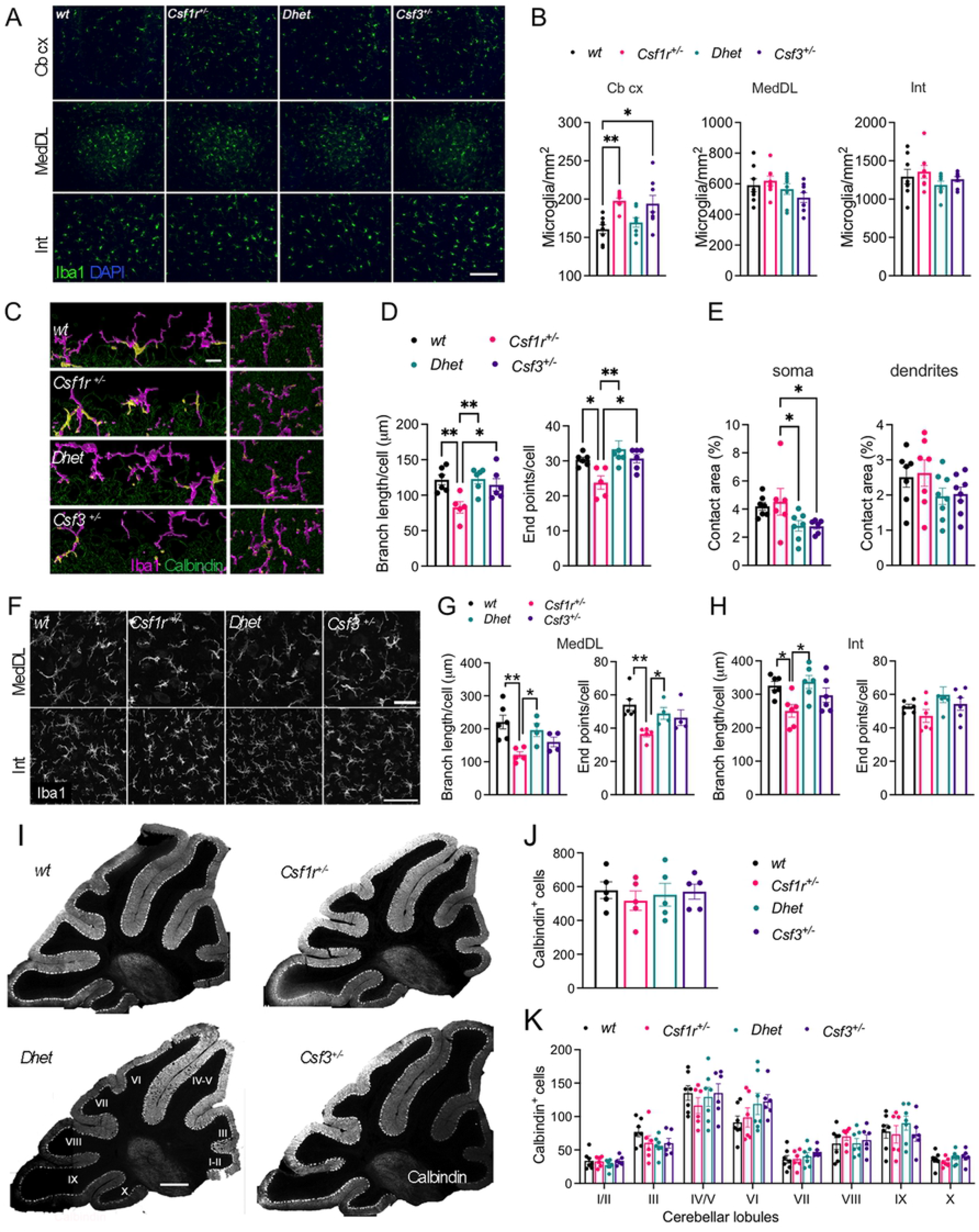
*Csf3* heterozygosity reduces microglial activation in the cerebellum of CRL mice. **(A)** Iba1^+^ cell densities (green) in the cerebellar cortex (Cb cx) and deep cerebellar nuclei (MedDL, dorsal protuberance of the medial cerebellar nucleus; Int, interposed nucleus) of 16-month-old female mice (scale bar, 100 μm). **(B)** Quantification of data in **(A)** (6-8 mice per genotype) (Bonferroni’s). **(C-E)** Imaging (**C** -left panels) and quantitation **(E)** of microglia contacts with the Purkinje cell somas and morphology (**C** -right panels) and morphometry **(D)** of microglia (red) in the cerebellar cortex of 16-month-old female mice (4-5 mice per genotype) (Dunn’s) (scale bar, 15 μm). **(F-H)** Morphology **(F)** and morphometry **(G, H)** of microglia in the deep cerebellar nuclei of 16-month-old female mice (scale bars, 50 μm and 70 μm respectively) (4-5 mice per genotype) (Dunn’s, and Tukey’s). **(I)** Representative images of Calbindin^+^ Purkinje cells (PC) distributed in the cerebellar lobules **(J)** Quantification of the total number PC per section. **(K)** Quantification of the number of Calbindin^+^ PCs in each lobule (4-5 mice per genotype) (scale bar, 500 μm). Means ± SEM, significantly different changes are marked by asterisks. *, p < 0.05; **, p < 0.01. Data underlying this figure can be found in Table S1.

Deletion of *Csf1* in the Nestin^+^ neural lineage, resulting in CSF1R signaling deficiency in the cerebellum, was associated, not only with alterations of cerebellar microglia homeostasis, but also with decreased Purkinje cell (PC) numbers (25). Furthermore, a reduction of PC number was also documented in *Csf1^op/op^* mice with global *Csf1* deficiency (26). These findings, together with our observation that G-CSF contributes to the activation of cerebellar microglia, prompted us to explore how *Csf1r* and/or *Csf3* heterozygosities impact PC number (Fig. 4 I-K). Neither the total numbers of Calbindin^+^ PC cells (Fig. 4J) nor their distribution in each lobule of the cerebellar cortex (Fig. 4K) were significantly different in mice carrying single or compound mutations in *Csf1r* and *Csf3*.

Together, these data indicate that, although *Csf1r* heterozygosity does not cause the loss of Purkinje cells, it promotes the activation of cerebellar microglia in a G-CSF-dependent manner.

### *Csf3* heterozygosity prevents deficits in social novelty

Aside from its role in motor coordination, the cerebellum is involved in regulation of aspects of social behavior (27). Both direct genetic disruption of Purkinje cell activity (28) and disruption of cerebellar microglia homeostasis in mice with neural-lineage specific deletion of *Csf1* (25) cause autistic-like behavior manifested as a loss of social novelty preference. This prompted us to investigate how decreased CSF-1R and/or G-CSF signaling impact social behavior. Using the three-chamber sociability paradigm for social preference and social novelty (Fig. 5A) we found that all mutant mouse groups exhibited normal social preference spending significantly more time interacting with another mouse than with an object (Fig. 5B). In contrast, *Csf1r^+/-^* mice showed a clear loss of social novelty, failing to preferentially interact with the novel mouse in comparison to the familiar mouse. This phenotype was prevented by monoallelic targeting of *Csf3* (Fig. 5C). These data suggest that G-CSF may contribute to the development of social interaction deficits in *Csf1r^+/-^* mice by impairing cerebellar function.

**Figure 5.**
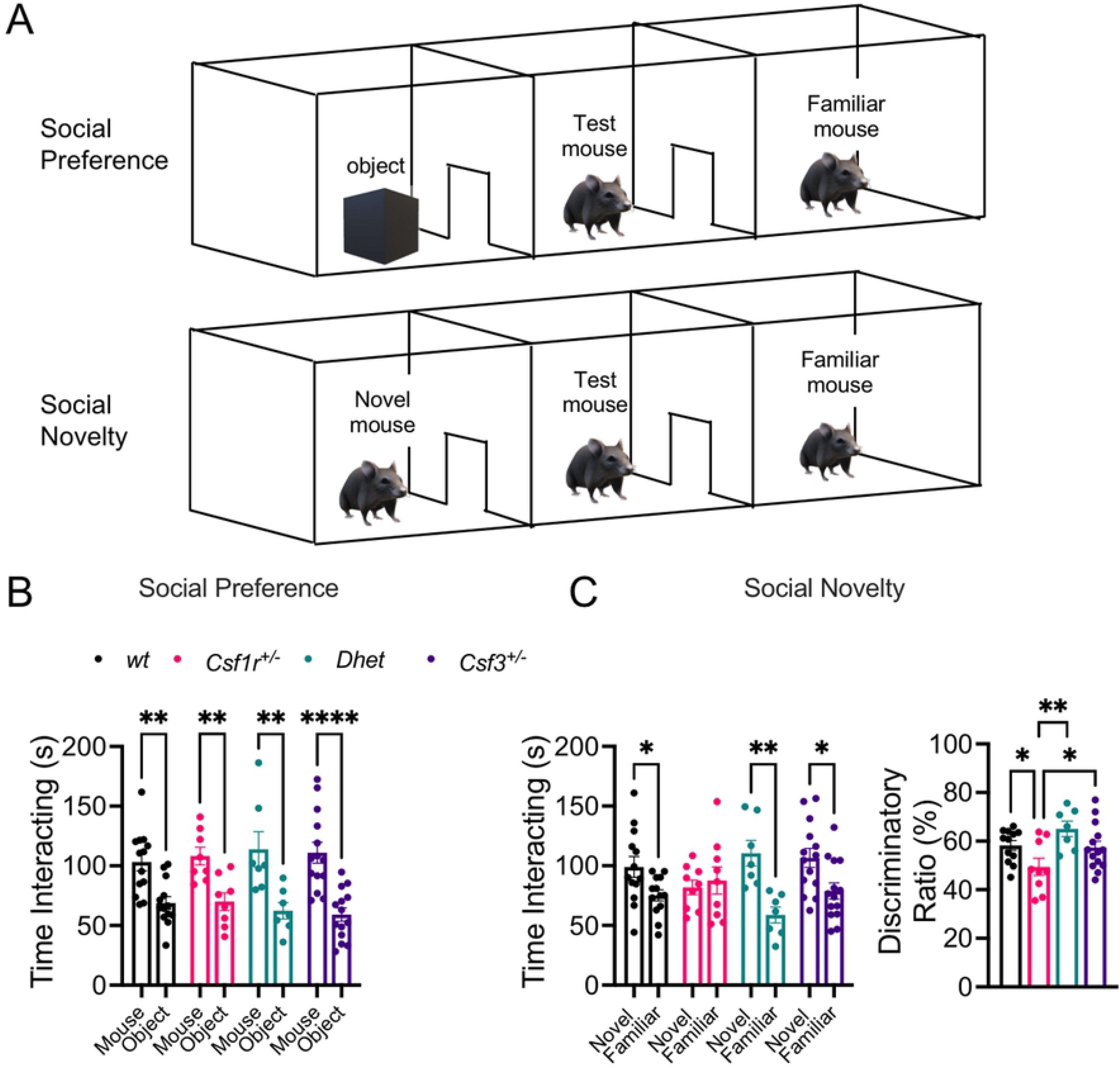
Evaluation of social interaction in the three-chamber sociability paradigm. **(A)** Schematic of the testing apparatus. Assessment of social preference **(B)** and social novelty **(C)** in the three chamber sociability paradigm. Combined female and male data. Means ± SEM, significantly different changes are marked by asterisks. *, p < 0.05; ****, p < 0.0001 (Holm-Sidak’s). Data underlying this figure can be found in Table S1.

### *Csf3* heterozygosity prevents the development of electrophysiological alterations in the deep cerebellar nuclei of *Csf1r^+/-^* mice

The attenuation of microglia density and activation in the cerebellar regions analyzed (Fig. 4) together with the correction of the cerebellum-dependent behaviors by *Csf3* heterozygosity (Fig. 1O and Fig. 5) suggested that G-CSF may play a role in microglia-mediated alteration of cerebellar physiology. To test this hypothesis, we analyzed the firing properties of PC and DCN. *In vivo* single cell unit recording of PCs (Fig. 6 A-E) and DCN cells (Fig. 6 F-J), revealed a differential effect of *Csf1r* heterozygosity. Electrophysiological recordings of the activity of Purkinje cells in awake, head-restrained mice revealed no difference in average firing rate (FR), predominant FR and inter-spike interval coefficient of variation (ISI CV) between *wt* and *Csf1r^+/-^* mice (Fig. 6 A- E). In contrast, recordings in the DCN revealed a significant decrease in predominant FR in *Csf1r^+/-^* mice that was normalized when one allele of *Csf3* was removed (Fig. 6 F-J). These data indicate that G-CSF mediates the disruption of DCN firing properties in CRL mice.

**Figure 6.**
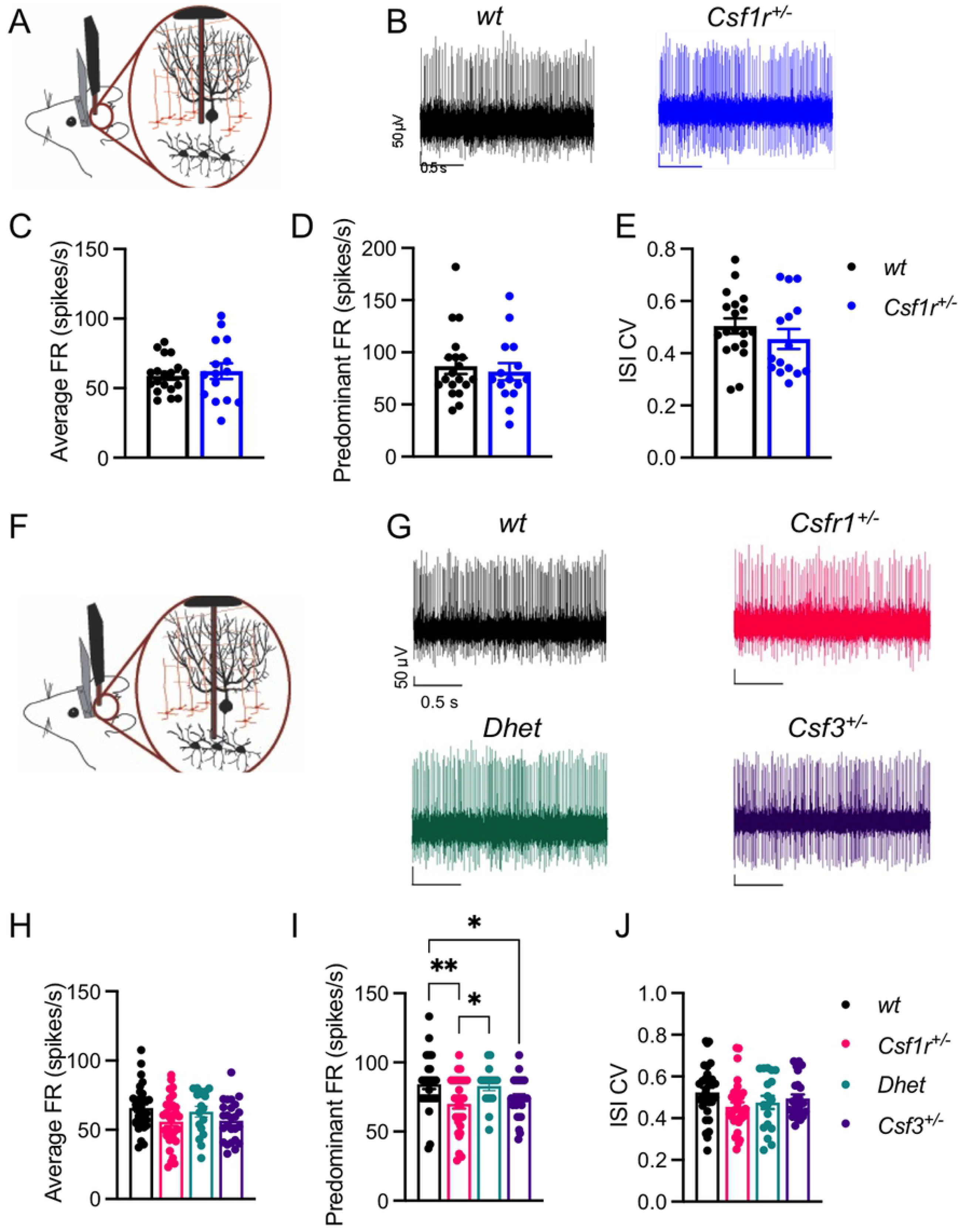
*Csf3* heterozygosity rescues the altered firing of deep cerebellar nuclei (DCN) cells in CRL mice. **(A)** Schematic of awake head-restrained *in vivo* single unit electrophysiological recording of cerebellar Purkinje cell (PC) activity. **(B)** Example recordings of PCs from *wt* (top left) and *Csf1r^+/-^* mice (top right). **(C-E)** Quantification of average firing rate (FR) **(C)**, predominant FR **(D)** and inter-spike interval coefficient of variation (ISI CV) **(E)** of sorted single units from wt (n = 19 cells, 3 mice) and *Csf1r^+/-^* (15 cells, 4 mice). **(F)** Schematic of *in vivo* single unit electrophysiological recording of DCN cell activity. **(G)** Example recordings of DCN cells from *wt* (top left), *Csf1r^+/-^* (top right), *Dhet* (bottom left) and *Csf3^+/-^* mice (bottom right). **(H-J)** Quantification of FR **(H)**, predominant FR **(I)** and ISI CV **(J)** of sorted single units/cells from *wt* (n = 32 cells, 5 mice), *Csf1r^+/-^* (31 cells, 4 mice), *Dhet* (17 cells, 3 mice) and *Csf3^+/-^* (25 cells, 4 mice). Means ± SEM, significantly different changes are marked by asterisks. *, p < 0.05; **, p < 0.01 (Fisher’s). Data underlying this figure can be found in Table S1.

### *Csf3* heterozygosity prevents the excessive elimination of glutamatergic synapses in the deep cerebellar nuclei of *Csf1r^+/-^* mice

Single cell transcriptome profiling experiments of mouse cerebelli detect *Csf3r* transcripts in microglia while the expression in neural lineage cells, including Purkinje cells, is sporadic, at best (29) (databases available at: https://singlecell.broadinstitute.org/single_cell/study/SCP795/a-transcriptomic-atlas-of-themouse-cerebellum?genes=Csf3r#study-visualize). Microglia play a pivotal role in remodeling neuronal networks by pruning or eliminating synapses during development and in adult life (30, 31). To investigate whether G-CSF-activated microglia may contribute to aberrant synapse pruning in the DCN of *Csf1r^+/-^* mice, we quantified the colocalization of microglia with synaptic markers of glutamatergic (VGLUT2^+^) and of GABAergic (GAD67^+^) neurons, the two prevailing neuronal populations in the DCN (32). While the percentage of GAD67^+^ puncta colocalized with Iba1^+^ microglia cells was comparable among all genotypes (Fig. 7 A, B), the percentage of VGLUT2^+^ puncta colocalized within Iba1^+^ microglial cells was significantly higher in the DCN of *Csf1r^+/-^* mice and was normalized by *Csf3* heterozygosity (Fig. 7 A, C). Three-dimensional reconstruction, revealed the presence of VGLUT2^+^ puncta in both the branches and the cell body of single microglial cells (Fig. 7A, lower panels), confirming the engulfment of synaptic material.

**Figure 7.**
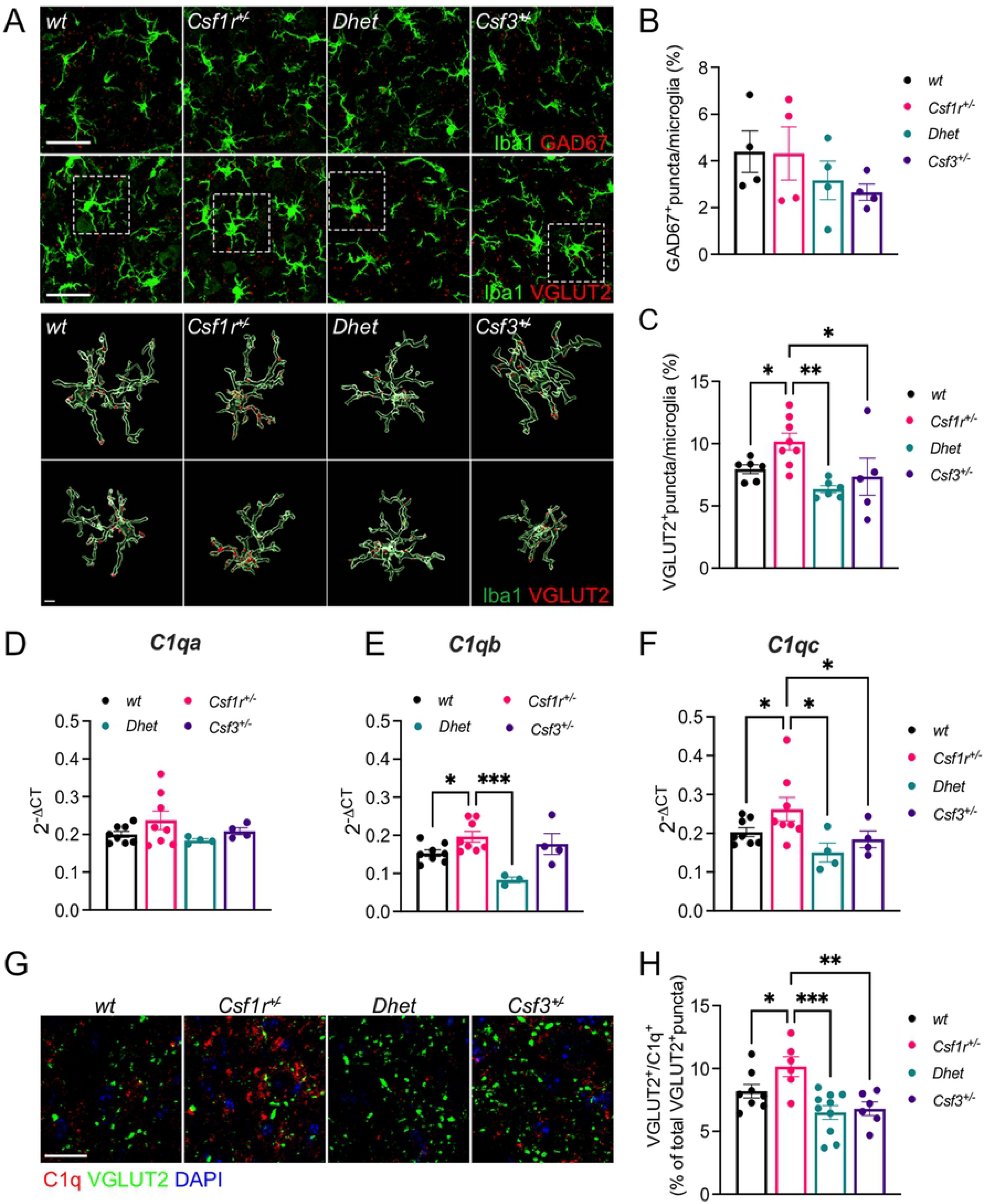
G-CSF mediates excessive complement-mediated engulfment of DCN glutamatergic synapses by microglia in CRL mice. **(A)** Upper panels, immunofluorescence staining showing the colocalization of GAD67^+^ and VGLUT2^+^ (red) with microglia (green) in the DCN of 16-month-old female mice. Lower panels, 3D reconstruction with surface rendering showing VGLUT2^+^ puncta inside selected individual microglia shown in their original position (top) and rotated 180 degrees along the z axis (bottom). Scale bars: 30 μm (upper panels), 5 μm (lower panels). **(B-C)** Quantification of the percentage of GAD67^+^ and VGLUT2^+^ puncta colocalized with Iba1^+^ cells (4-8mice per genotype; Newman-Keuls). **(D-F)** Quantification of the expression of transcripts of C1q genes *C1qa*, *C1qb*, and *C1qc* in the cerebella of 16-month-old female mice (4-8 mice per genotype; two-stage linear step-up procedure of Benjamini, Krieger and Yekutieli). **(G, H)** Co-localization of VGLUT2^+^ puncta (green) and C1q (red) in the DCN of 16-month-old female mice (6-10 sections per genotype; Holm-Sidak’s). Means ± SEM, significantly different changes are marked by asterisks. *, p < .05; **, p < 0.01, ***, p < 0.001. Scale bar in **(G)**, 20 μm. Data underlying this figure can be found in Table S1.

### *Csf3* heterozygosity prevents the overexpression of C1q genes and attenuates C1q deposition on glutamatergic synapses in *Csf1r^+/-^* mice

The complement cascade of the innate immune system has emerged as an important mediator of synapse pruning during both brain development and disease (33, 34). The C1q factor, comprising 6 C1qA, 6 C1qB and 6 C1qC chains, is the initiating protein of the classical complement cascade and was reported to associate with synapses to promote their removal by microglia (34). These findings prompted us to analyze the expression of C1q genes in the cerebella of the mouse models analyzed and the extent of C1q deposition on glutamatergic synapses of DCN. Quantitative RT-PCR revealed significantly increased expression of *C1qb* and *C1qc* transcripts in *Csf1r^+/-^* mice, that was normalized by monoallelic targeting of *Csf3* (Fig. 7 D-F). The increased expression of *C1q* genes in *Csf1r^+/-^* mice was associated with increased C1q deposition on glutamatergic synapses, evidenced by the increased colocalization of C1q with VGLUT2^+^ puncta (Fig. 7 G, H) and was normalized by monoallelic deletion of *Csf3.* In contrast, the expression of genes encoding other components of the complement cascade (*C3),* complement receptors *(Itgam, Itgax, Itgb2*), of. neuronal proteins that mediate the synaptic deposition of C1q (*Nptx1, Nptx2)* (35) and of *Trem2*, a microglial receptor involved in synapse removal (36) was unchanged (Supplementary Fig. 1). Overall, these data, together with the identification of microglia as the dominant source of C1q in the mouse brain (37), indicate that the elevated G-CSF signaling in CRL mice causes an excessive removal of glutamatergic synapses in the DCN through the activation of microglia.

## Discussion

In a previous study, we demonstrated that *Csf1r^+/-^* mice reproduced the hallmark features of CRL patients (9). The cognitive, emotional and motor deficits were accompanied by histological alterations including elevated brain microglial density, callosal demyelination, cortical neuronal loss, and callosal axonal spheroids. These phenotypes were associated with increased brain expression of *Csf2,* encoding GM-CSF, and of *Csf3,* encoding the G-CSF (9). Subsequent studies in autopsied brain tissue of CRL patients showed increased expression of *CSF2* (10) and provided evidence of dysregulated G-CSF signaling (6, 7) suggesting an important role for these factors in CRL. Since the CSF-1R, as well as the receptors for both G- and GM-CSF, are predominantly expressed in microglia and regulate their activation (reviewed in (11)), it was suggested that CRL could be a primary microgliopathy. Indeed, monoallelic deletion of *Csf1r* in the microglial lineage recapitulated the phenotype observed in *Csf1r^+/-^* mice, indicating that CRL is a primary microgliopathy (8). Targeting *Csf2*, that encodes a microglial mitogen, rescued some behavioral defects (spatial memory, depression and olfactory) and the histological alterations observed in the forebrain of *Csf1r^+/-^* mice (microgliosis, callosal demyelination, decreased callosal volume) but did not attenuate the elevated microglial density in the cerebellum (10). Furthermore, the expression of *CSF3* was also elevated in post-mortem CRL brains (Fig. 1A). While G-CSF is not a microglial mitogen (38), its administration was reported induce the expansion of microglia *in vivo* (39), to activate a Cathepsin S-CX3CR1-inducible NOS pathway in microglia and to induce the production of factors that promote neuronal excitability (15). Therefore, we explored whether G-CSF may contribute to CRL pathogenesis.

A hallmark feature of CRL is the loss of callosal white matter (reviewed in (1, 5)). The primary function of the corpus callosum is to integrate the information by joining both cerebral hemispheres to process motor, sensory, and cognitive signals and disruption of myelination could potentially impact all these functions. Indeed, a study employing advanced MRI techniques revealed altered functional connectivity between the cerebral hemispheres in CRL patients and highlighted an association between their symptoms and the disconnection of the two cerebral hemispheres, due to the loss of connection fibers in the corpus callosum (40). Furthermore, studies in both autopsied tissue from CRL patients and the mouse model suggest that microglial activation contributes to the loss of callosal white matter (8, 10, 41, 42). Intriguingly, although targeting *Csf3* reduced the density of microglia in the corpus callosum, in contrast to *Csf2* reduction (10), it did not prevent their activation and clustering, nor did it prevent demyelination. In addition, targeting *Csf3* did not prevent the loss of layer V neurons, a population that is uniquely dependent on trophic support from microglia (43). Together, these data indicate that G-CSF does not play a major role in promoting demyelination or cortical neurodegeneration in the mouse model of CRL.

The memory deficits of *Csf1r^+/-^* mice developed independently of the level *Csf3* expression. Furthermore, consistent with a previously published study in *Csf3^-/-^* mice (44), we show that monoallelic deletion of *Csf3* was sufficient to cause cognitive dysfunctions in wild type mice, a phenotype that could be related to the reduction of its neurogenic actions in the dorsal hippocampus. Nevertheless, monoallelic targeting of *Csf3* reduced microglial activation in the ventral hippocampus and ameliorated anxiety-like behavior in female mice. The factors contributing to the differential effects of G-CSF in different areas of the hippocampus remain to be elucidated. However, the rescue of two cerebellum-dependent functions, i.e. the motor and social interaction deficits (27, 45) by *Csf3* heterozygosity suggested that G-CSF mediates cerebellar dysfunction in CRL mice, a hypothesis that is supported by the histological data showing attenuation of microglial activation in all areas of the cerebellum following monoallelic targeting of *Csf3* in *Csf1r^+/-^* mice. *Csf3* heterozygosity also decreased the elimination of glutamatergic synapses and restored electrophysiological activity in the deep cerebellar nuclei. Furthermore, these effects of *Csf3* heterozygosity were associated with normalization of the expression of *C1q* genes and of C1q deposition on glutamatergic synapses. Since *Csf3r* expression within the brain is largely restricted to microglia (46) (databases available at https://portals.broad institute.org/single_cell/study/aging-mouse-brain, http://dropviz.org/), we conclude that G-CSF mediated activation of microglia in specific brain regions promotes the development of anxiety-like behavior, motor coordination and social interaction deficits in *Csf1r^+/-^* mice.

Females tend to be more severely affected by CRL and exhibit a higher prevalence of gait disorders (5). Furthermore, ataxia and cerebellar involvement have also been reported predominantly in female CRL patients (13 out of 15 documented cases) (42, 45, 47–55). These findings raise the possibility that estrogens and/or androgens might specifically regulate subpopulations of microglia such as the cerebellar microglia, or those interacting with motor neurons. Remarkably, targeting *Csf3* selectively rescued the motor coordination deficits of female mice, while it also tended to worsen motor function in males (Fig. 1 P). This finding is not unique to the CRL mouse model. Administration of G-CSF has also produced gender-specific effects in both preclinical and clinical trials for amyotrophic lateral sclerosis, providing protection in males by attenuating inflammation and exacerbating the loss of motor function in females (56–58). These data suggest an interaction of G-CSF with gender-specific factors, likely hormonal, in the control of neuroinflammation, an aspect that deserves further exploration.

In conclusion, this study identifies elevated G-CSF as the main factor driving anxiety and cerebellar dysfunctions contributing to the motor coordination and social preference deficits in CRL. Apart from their overlapping contributions to the motor coordination deficit, the effects of elevated G-CSF and GM-CSF in CRL are non-redundant (10) (Table 1). Thus, our studies point to G-CSF as an additional potential therapeutic target in CRL.

## Materials and methods

### Mouse strains, breeding, and maintenance

Experiments were performed on adult C57BL/6J mice (RRID: IMSR JAX:000664) of the indicated ages and genders. The generation, maintenance and genotyping of *Csf1r^+/-^* mice was described previously (59). *Csf3^+/-^* mice on a mixed C57BL/6 x 129/Ola genetic background (60) were obtained from Jackson Laboratories and genotyped by PCR utilizing the following primers: Csf3- Fw (5’- GCACCCTCAGTATCCTTCCA-3’), Csf3-Rev (5’- GCTAGAGCAGCCACTCAGG -3’) and Csf3-Neo (5’-GCTATCAGGACATAGCGTTGG-3’) specific for the neomycin resistance gene. Both lines were backcrossed for more than 10 generations onto the C57BL6/J background. Cohorts were developed from the progeny of matings of *Csf1r^+/-^* to *Csf3^+/-^* mice, randomized with respect to the litter of origin and maintained on a breeder (PicoLab Rodent Diet 20 5058) rather than a maintenance diet, in order to accelerate symptom development (5). The age and sex of mice used in each experiment are indicated in the figures. All *in vivo* experiments were performed in accordance with the National Institutes of Health regulations on the care and use of experimental animals and approved by the Institutional Animal Care and Use Committees of Albert Einstein College of Medicine and Hunter College.

### Behavioral studies

Male and female mice were tested sequentially for memory, anxiety, motor coordination and social interaction. A separate cohort was developed for active place avoidance. All the experiments were conducted by a blinded experimenter during the light cycle. The animals were allowed to acclimate to the behavior room for one hour before the beginning of each experiment.

For each experimental paradigm, mice were randomized and balanced to avoid unwanted effects of confounding factors.

### Cognitive flexibility

Cognitive flexibility was evaluated in the active place avoidance paradigm at 7 months of age (21). The apparatus consisted of a circular 40 cm diameter platform rotating clockwise at 1 rpm. A camera placed above the apparatus recorded the mouse location during each stage of the experiment. The apparatus was controlled by PC-based software (Tracker, Bio-Signal Group Corp., Brooklyn, NY) that tracked the mouse position and delivered a foot shock (500 ms, 60 Hz,1.2 2 mA) every time the mouse was inside a 60° stationary shock zone that could be identified by visual cues within the room. The number of entrances into the shock zone was recorded throughout the duration of the experiment.

The task included four steps:

1. Habituation. Mice were allowed to freely explore the apparatus in the absence of shock for 30 minutes.
2. Training. For three consecutive days, mice were placed on the apparatus with the shock turned on. Each day, mice were trained with a single 30-minute trial to avoid one shock zone. The location of the shock zone was constant across trials.
3. Long-term memory test. Mice were returned to the apparatus three days after the last training day with the shock turned on. During this 10-minute trial, the shock zone remained in the same location as the previous training trials.
4. Cognitive flexibility test. Two hours after the long-term memory test, the location of the shock zone was moved to the opposite side of the arena, which is where mice primarily spent their time on the previous trials. The number of entrances into the new shock zone was recorded over a 20 min period.

### Spatial memory

Short-term memory. Mice were assessed at 11.5 months of age for short-term spatial memory in the two-stage version of the Y-maze (61). In the first training stage, each mouse was introduced into the Y-maze and allowed to explore two of the three arms of the apparatus. In the second testing stage, conducted one hour later, the remaining arm was opened and the mouse returned to the apparatus to freely explore all the three arms. Internal visual cues were placed inside each arm as referential tools to explore the maze. The number of arm entries into each arm was tracked and recorded by Any-maze (Stoelting). The positions of the three arms were randomized within each genotype.

Long-term memory. Long-term spatial recognition memory was evaluated in 11.5-old month mice using the object placement test. Each mouse was allowed to interact with two identical objects placed 10 cm apart parallel to one of the walls of a 40 cm x 40 cm chamber for 10 minutes (training). After 24 hours, one of the objects was displaced into a novel position (15 cm distant, 90°angled) and the mouse returned to explore the objects for 10 minutes (testing). Visual cues were affixed to the walls of the chamber to assist orientation within the arena. Time interacting with the objects was tracked by a blinded experimenter.

### Anxiety-like behavior

Anxiety was measured by using the elevated zero maze (Ugo Basile Instruments) at the age of 12 months. The apparatus consisted of an elevated ring-shaped apparatus (diameter 50 cm, width 5 cm) including two opposite open zones, and two opposite enclosed zones. Each mouse was allowed to explore the apparatus for 3 minutes. Cumulative time spent in the open zones was tracked by ANY-maze software (ANY-maze, Stoelting), and utilized as a measure inversely related to anxiety (62).

### Motor coordination

Motor coordination was tested in the balance beam test (63). The balance beam consisted of a 1-meter-long wooden beam (1.6 cm in diameter) elevated 50 cm above the floor. Each mouse was positioned at one end of the beam and encouraged to cross the beam. The presence of palatable food placed at the opposite end was used as reinforcement to accomplish the task. The number of slips tracked by the experimenter were used as measure of motor coordination.

### Social interaction

Sociability was tested in 10-18-month-old mice using the three-chamber sociability test (25). This paradigm is based on the natural tendency of mice to preferentially interact with other mice rather than with an inanimate object (social preference), and with a novel mouse rather than with a familiar mouse (social novelty). The three-chamber apparatus consisted of a white Plexiglas box (60 x 40 x 15 cm) divided into three chambers (20 x 40 x 15 cm) by two transparent Plexiglas walls (40 x 15 cm). Entry from the middle chamber to each lateral chamber was made accessible by removable sliding doors (9 x 5.5 cm) (Fig. 5a). The experiment consisted of three 10-minute consecutive stages: habituation, social preference test and social novelty test. Before the start of each stage, the experimental mouse was confined to the middle chamber by the dividing doors. In the habituation, each mouse was allowed to explore the whole empty apparatus. In the social preference test, the mouse was exposed to an object (a plastic black cube) and to another mouse (familiar mouse). In the social novelty test, the object was replaced by another mouse (novel mouse), and the experimental mouse allowed to explore the apparatus and interact with the mice. The object, the familiar mouse and the novel mouse were placed under wire mesh pen cups (11.5 cm high, 9.5 cm in diameter) when introduced into the apparatus. The time interacting with each object or mouse was recorded by ANY-maze video tracking system (ANY-maze, Stoelting).

### Human studies

Frozen brain tissue blocks containing periventricular white and grey matter were obtained from the Mayo Clinic Brain Bank. Consent for autopsy was obtained from the legal next-of-kin. Information on the CRL patients harboring CSF1R mutations and control cases included in this study is summarized in Table S2. Upon removal from the skull according to standard autopsy pathology practices, the brain was divided in the mid-sagittal plane. Half was fixed in 10% neutral buffered formalin, and half was frozen in a -80°C freezer, face down to avoid distortion. The frozen brain was shipped on dry ice to the Neuropathology Laboratory at Mayo Clinic, where it was stored in a -80°C freezer. Frozen tissue was partially thawed before dissection and slabbed in a coronal plane at about 1-cm thickness. Regions of interest were dissected from the frozen slabs and placed in microcentrifuge tubes before being shipped to the research laboratory on dry ice. At all steps, the fresh and frozen tissue was handled with Universal Precautions.

### Gene expression in CRL patients and mouse brains

RNA was isolated from the gray matter of 5 CRL patients and 5 control patients (see Supplemental Table 1) using Trizol and cDNA was prepared using a Super Script III First Strand Synthesis kit (Invitrogen, Carlsbad, CA). Real time PCR was performed using the PrimePCR CSF3 assay qHsaCED0043218 from BIO-RAD. Human *RPL13* (Fw: 5’-AGCCTACAAGAAAGTTTGCCTAT-3’; Rev: 5’-TCTTCTTCCGGTAGTGGATCTTGGC-3’) was used for normalization. Average values from two different blocks of tissue per patient, were used to construct the figure.

For mouse studies, the RNA was extracted from the anterior motor cortex, corpus callosum and cerebellum of 6-month-old mice as described (9), reverse-transcribed as described above and the qPCR was carried out utilizing SYBR Green in an Eppendorf Realplex II thermocycler. Beta actin was used as a housekeeping gene control. The primers for mouse genes used were as follows: *Csf3* (Fw: 5’-GAGCAGTTGTGTGCCACCTA-3’; Rev: 5’- GCTTAGGCACTGTGT CTGCTG-3’), *C1qa* (Fw: 5’-GGATGGGGCTCCAGGAAATC- 3’; Rev: 5’- CTGATA TTGCCTGGATTGCC- 3’), *C1qb* (Fw: 5’-TGGCTCTGATGGCCAACCAG-3’; Rev: 5’-GACTTTCTGTGTAGCCCCGT-3’), *C1qc* (Fw: 5’-AGGACGGGCATGATGGACTC- 3’; Rev: 5’- TGAATACCGACTGGTGCTTC-3’), *C3* (Fw: 5’-CGCAACGAACAGGTGGAGATCA- 3’; Rev: 5’- CTGGAAGTAGCGATTCTTGGCG-3’), *Itgam* (Fw: 5’-CTGAGACTGGAGGCAACCAT-3’; Rev: 5’- GATATCTCCTTCGCGCAGAC-3’), *Itgb2* (Fw: 5’-CCCAGGAATGCACCAAGTACA-3’; Rev: 5’- CAGTGAAGTTCAGCTTCTGGCA- 3’), *Itgax* (Fw: 5’- CTGGATAGCCTTTCTTCTGCTG- 3’; Rev: 5’- GCACACTGTGTCCGAACTCA-3’), *Nptx1* (Fw: 5’-ATCACCCCATCAAACCACAG-3’; Rev: 5’ CGATGACATTGCCAGAGAGA-3’), *Nptx2* (Fw: 5’-CGGAGCTGGAAGATGAGAAG-3’; Rev: 5’- GGAAGGGACACTTTGAATGC-3’), *Hprt* (Fw: 5’-CAAACTTTGCTTTCCCTGGT-3’; Rev: CAAGGGCATATCCAACAACA), *Actb* (Fw: 5’- AGAGGGAAATCGTGCGTGAC-3’; Rev: 5’- CAATAGTGATGACCTGGCCGT-3’).

### Immunofluorescence staining and data analysis

Immunostaining was performed in brain slices prepared as described previously (8). Brain sections were incubated with primary antibodies overnight at 4°C. The primary antibodies used in the study included: Iba1 (1:500) (rabbit IgG; Wako Chemicals RRID: AB_839504 or goat IgG; Abcam RRID:AB_10972670); Calbindin, (1: 500) (mouse IgG, Abcam RRID:AB_1658451); NeuN (1:500) (mouse IgG, Millipore RRID:AB_2149209); GAD67 (1:500) (mouse IgG, Millipore RRID:AB_94905); VGLUT2 (1:500) (polyclonal guinea pig antiserum, Synaptic Systems RRID:AB_887884). Following incubation with primary antibodies, the sections were incubated with secondary antibodies conjugated to either Alexa 488, Alexa 594, or Alexa 647 (1:500) (Life Technologies) for 1 hour at room temperature. Fluoromyelin staining for myelin (1:350, 30 minutes) was performed according to the manufacturer’s (Molecular Probes, Inc.) instructions.

For C1q staining, slices were blocked with 5% bovine serum albumin (BSA) and 0.2% Triton X-100 solution for 1 h and incubated with primary antibody overnight (1:500) (rabbit IgG, Abcam RRID:AB_2732849). After washing, the secondary antibody was applied and incubation continued for 2 hours at room temperature (64). Sections were mounted on SuperFrost Plus slides (Thermofisher) using Prolong antifade mountant with DAPI (Thermofisher). Images were captured using a Nikon Eclipse TE300 fluorescence microscope with NIS Elements D4.10.01 software. Cell number quantification was performed manually. Quantification of fluorescent areas was performed using ImageJ.

For confocal microscopy, microscope Z series stacks were obtained by a Leica SP8 Confocal microscope at ×40 magnification with a 0.40 μm interval between stacks. Images were cropped and adjusted for brightness, contrast and color balance using Adobe Photoshop CC. For analysis of synapse engulfment, Imaris software (Oxford Instruments Group) was used to analyze colocalization and to generate 3D reconstructions and surface renderings (65).

Morphometric analysis of microglia (number of end points and length of cell processes) was performed on maximum intensity projections of tissue sections using FIJI as previously described (10, 66). The extent of microglia-Purkinje cell contacts was examined in confocal 3D surface rendering images using Imaris as described (67).

### *In vivo* electrophysiology

*In vivo* single unit recording in the cerebellum was performed in awake head-restrained mice. Mice were anesthetized with isoflurane (5% induction, 2% maintenance), the head shaved, wiped with ethanol and betadine and the scalp reflected to reveal the skull. The skull was lightly scraped and cleaned with ethanol. Recording coordinates were then marked by lightly drilling over the interparietal bone which overlies the cerebellum and touching the drilled sites with a marker pen. The skull was then covered with OptiBond (Kerr Corporation, Brea, CA, USA) and cured with ultraviolet light. A titanium bracket was subsequently fixed onto the skull with Charisma (Kulzer GmbH, Germany) just anterior to the lambdoid suture, enabling later access to the cerebellum, and covered with dental cement (M& Dental Supply, Jamaica, NY, USA). A recording chamber was simultaneously created over the interparietal bone and once the cement dried, the chamber was covered with Kwik-Sil silicone elastomer (World Precision Instruments, Sarasota, FL, USA).

The mice were monitored post-surgery for 1 week before neural recordings. During this time, the mice were acclimated to head-restraint using screws to immobilize the previously implanted bracket. This was done for 0.5-1 h per day. Twenty-four hours prior to recording, a craniotomy was created by removing the Kwik-Sil covering the recording chamber and drilling at coordinates previously marked over the interparietal bone. Purkinje cells were recorded at AP: -6.00 mm, ML: 0 mm and AP: -7.0 mm, ML: 0mm; and deep cerebellar nuclei at AP: - 6.2 mm, ML: ± 1.5 mm. The recording chamber was recovered with Kwik-Sil and the mouse was allowed to recover.

For single unit recordings, the mouse was head restrained next to a stereotaxic apparatus on a padded flat air table using the head bracket and screws. The Kwik-Sil was removed from the recording chamber, revealing the craniotomies. A ground electrode was then placed into the recording chamber, a tungsten electrode (2-3 MΩ, Thomas Recording, Giessen, Germany) lowered under microscopic guidance into a craniotomy until it touched the surface of the brain/cerebellum, and the chamber filled with saline. In single unit recording, the electrode was further slowly advanced into the cerebellum until neuronal activity was detected but not more than 3 mm below the surface of the brain. Purkinje cell activity was identified by their characteristic firing rate, location and the brief pauses in firing following complex spikes. DCN cells were identified based on their location and firing. Cells were recorded for 2-5 mins. Neural signals were filtered at 20 kHz, amplified at 2000X on a custom amplifier, digitized at 20kHz with a National Instruments BNC-2110 (National Instruments, Austin, TX, USA) analog to digital converter into a PC and visualized with LabView (National Instruments, Austin, TX, USA). Waveforms of recorded single unit activity were sorted offline using Offline Sorter software (Plexon, Dallas, TX, USA) and analyzed using a custom LabView script to obtain the average firing rate, predominant firing rate and the interspike interval coefficient of variation (ISI CV).

### Statistical analyses

Statistical analyses were computed using GraphPad Prism 8 (GraphPad, La Jolla, CA). Data were checked for outliers using the Grubbs’ method. Gaussian distribution was evaluated using the Shapiro-Wilk normality test and the Kolmogorov-Smirnov test. The screened data were analyzed using the Student t-test, the Kruskal–Wallis test or by analysis of variance (one- or two-way ANOVA). When significant effects of the independent variables were detected, single differences between or within genotypes were analyzed by post-hoc multiple comparison tests (Dunnett’s, Bonferroni, Tukey’s, Fisher’s LSD, and the two-stage linear step-up procedure of Benjamini, Krieger and Yekutieli as indicated in the figure legends). The level of significance was set at p < 0.05. For those comparisons in which no statistical significance is indicated in the figure panels, the p value was > 0.05. Data are presented as mean ± SEM. Sample sizes for each experiment are indicated in the figure legends.

### Data Availability

The data sets used and analyzed during the current study are available from the corresponding author on reasonable request.

## Acknowledgements

The authors thank Hillary Guzik, Andrea Briceno and Dr Vera Des-Marais of the Einstein Analytical Imaging Facility for help with imaging and histomorphometry, Dr. Daniel Wilton of Dr. Beth Stevens laboratory for sharing their protocol for C1q staining and Christopher Fernandes and Jude Oppong-Asare for technical assistance.

## Funding disclosure

This work was supported by grants from the National Institutes of Health: Grant R01NS091519 (to E. R. S.), R01NS105470 (to K.K.), R21MH114182 (N.S.B), U54 HD090260 (support for the Rose F. Kennedy IDDRC), the P30CA013330 NCI Cancer Center Grant and a gift from David and Ruth Levine. ZKW is partially supported by the NIH/NIA and NIH/NINDS (1U19AG063911, FAIN: U19AG063911), Mayo Clinic Center for Regenerative Medicine, Mayo Clinic in Florida Focused Research Team Program, the gifts from The Sol Goldman Charitable Trust, and the Donald G. and Jodi P. Heeringa Family, the Haworth Family Professorship in Neurodegenerative Diseases fund, and The Albertson Parkinson’s Research Foundation.

## Competing interests

The authors declare that they have no competing interests.

## Supporting information

Fig S1. Expression of genes involved in synapse removal in the cerebellum. The expression of genes encoding the C3 component of the complement cascade (*C3),* complement receptors (*Itgam, Itgax, Itgb2*), *Trem2*, and neuronal proteins that mediate the synaptic deposition of C1q (*Nptx1, Nptx2)* was analyzed by qPCR. Means ± SEM. Data underlying this figure can be found in Table S1.

Table S1. Underlying numerical data for Figures: 1A, 1B, 1D, 1E, 1F-P, 2B, 2C, 2E, 2F, 3B, 3D, 3E, 3J, 3L, 4B, 4D, 4E, 4G, 4H, 5B, 5C, 6C-E, 6H-J, 7B-D, 7H and S1.

Table S2. Summary of Subjects Analyzed for *CSF3* expression.

## Notes

### Competing Interest Statement

The authors have declared no competing interest.

